# Virtual spatial transcriptomics from histopathology enables prognostic and therapeutic response prediction in cancer

**DOI:** 10.64898/2026.08.04.742671

**Authors:** Shaoqing Jiao, Zhen Yuan, Dazhi Lu, Yunfei Xu, Yunwei Dong, Jiajie Peng

**Affiliations:** School of Software, Northwestern Polytechnical University, No.1 Dongxiang Road, Xi’an,710129,Shaanxi, China; AI for Science Interdisciplinary Research Center, School of Computer Science, Northwestern Polytechnical University, No.1 Dongxiang Road, Xi’an, 710129,Shaanxi, China; Key Laboratory of Big Data Storage and Management, Northwestern Polytechnical University, Ministry of Industry and Information Technology, No.1 Dongxiang Road, Xi’an,710129,Shaanxi, China; Department of General Surgery, Qilu Hospital of Shandong University, 107 Wenhuaxi Road, Jinan, 250012, Shandong, China

**Keywords:** Virtual spatial transcriptomics, Pathology foundation models, Gene expression prediction, Deep learning

## Abstract

Spatial transcriptomics reveals cellular heterogeneity, intercellular communication, and tissue organization, but its cost and limited accessibility restrict clinical use. Here, we present VISTA, a model that integrates multi-scale histological features and spatial context to infer spatial gene expression from H&E-stained tissue images. Across leave-one-section-out cross-validation and independent validation, VISTA robustly predicted thousands of genes and outperformed state-of-the-art methods. Beyond expression reconstruction, VISTA enabled clinically relevant downstream analyses. In TCGA breast cancer samples, it identified survival-associated genes, stratified prognostic risk groups, and revealed adverse tumor-associated spatial subtypes. In our in-house intrahepatic cholangiocarcinoma cohort, it preserved tumor–normal organization and identified CLDN4 and CYP3A4 as complementary spatial biomarkers. In HER2+ breast cancer, it predicted pathological response to neoadjuvant trastuzumab-based therapy and linked response-associated regions to immune and cytokine-related programs. These results support virtual spatial transcriptomics from routine histopathology for oncology applications.

## 1 Introduction

The advent of spatial transcriptomics (ST) has enabled comprehensive analysis of cell-type composition and spatial distribution by linking transcriptomic profiles to their spatial locations [1, 2]. Leveraging this spatial information, ST has provided new insights into immune cell responses and features of the tumor microenvironment, improving our understanding of tumorigenesis, progression, and patient prognosis [3, 4]. Although recent ST technologies hold promise for new biological insights, the high cost limits their adoption in routine clinical practice [5]. In contrast, hematoxylin-and-eosin-stained (H&E) histopathology images are cost-effective and remain a diagnostic standard in cancer care [6]. Moreover, recent advances in deep learning have demonstrated that these images contain rich morphological information, enabling the identification of genetic mutations, bulk mRNA expression, and patient outcomes [7–9]. Given that spatial transcriptomics data are often paired with aligned H&E images, predicting spatial gene expression directly from H&E slides offers an opportunity to extend these insights at scale [10, 11].

Deep learning–based methods have been developed to predict spatial gene expression from H&E images, yet their predictive performance remains limited. Existing methods can be broadly categorized into two types. One type of method predicts gene expression from H&E image patches using convolutional neural networks or Transformer [12] backbones to learn predictive representations. For example, Hist2ST [13] integrated a Transformer with graph neural networks to model spatial dependencies, followed by zero-inflated negative binomial modeling of gene expression values. Similarly, EGNv1 [14] employs a Vision Transformer backbone with exemplar learning to guide predictions. Several studies further explored richer local representations by incorporating graph-based and dynamic convolutional architectures. TCGN [15] applied convolution and Graph-Node co-embedding to characterize local spot-level spatial relationships, while THItoGene [16] combines dynamic convolution, capsule networks [17], and graph attention networks [18] to capture cross-regional associations and neighborhood interactions. However, these methods are typically trained in a task-specific manner on relatively limited spatial transcriptomics datasets [5]. As currently available spatial transcriptomics data remain small in scale, the learned representations may not fully capture rich histopathological features associated with gene expression. More recent methods, including DeepPT [8], FmH2ST [19], and MISO [20], have further leveraged pretrained image models to enhance feature extraction from histopathology images. For example, DeepPT employed a pretrained ResNet encoder to extract tile-level features, followed by a fully connected layer for spot-wise gene expression prediction. FmH2ST incorporated image foundation models with Transformer and graph neural network modules to enhance spot-level representations. MISO utilized H0-mini [21] for tile feature extraction and aggregated whole-slide information via attention-based multiple instance learning. Despite these advances, existing methods remain limited in capturing both fine-grained histological details and broader tissue-level context. In particular, tile-based representations often process fixed-size image patches independently, which can restrict the effective receptive field and weaken the modeling of cross-region morphological dependencies [22, 23]. Moreover, many methods do not explicitly adapt to spatial heterogeneity in cell density and tissue architecture, limiting their capacity to represent biologically meaningful variation across tissue regions. These limitations highlight the need for a method that integrates multiscale histological features while preserving fine-grained morphological information for spatially resolved gene expression prediction.

To address these challenges, we developed VISTA, a multi-scale and spatially aware model for predicting spatial gene expression from H&E images. VISTA leverages pretrained pathology foundation models to capture local morphological features and slide-level contextual information. A Global-to-Local Cross-Attention mechanism expands the receptive field beyond individual spots, while a Spatial-Position-Aware module with GatedFusion adaptively integrates spatial coordinates and visual features. Comprehensive benchmarking using leave-one-section-out cross-validation and independent validation demonstrated superior performance over existing state-of-the-art methods. When applied to TCGA breast cancer samples, VISTA identifies survival-associated genes, stratifies patients into prognostic risk groups, and reveals spatial subtypes driven by tumor-associated domains linked to adverse outcomes. In an independent intrahepatic cholangiocarcinoma cohort collected by our group, VISTA preserves tumor–normal spatial organization, recapitulates disease-relevant programs, and identifies CLDN4 and CYP3A4 as complementary spatial biomarkers. Finally, in a HER2+ breast cancer cohort treated with neoadjuvant trastuzumab-based therapy, VISTA predicts pathological response and links model-attributed regions to immune infiltration, cytokine signaling, and chemokine-mediated cell communication. These results establish VISTA as a scalable and cost-effective method for augmenting large-scale histopathology cohorts with in silico spatial transcriptomics and extracting clinically relevant spatial biomarkers for precision oncology.

## 2 Results

### 2.1 Overview of VISTA

We propose VISTA, a multi-scale and spatially aware model for predicting spatial gene expression from H&E slide images. VISTA integrates complementary feature representations across three different scales, enabling accurate prediction of spatial gene expression. For each tissue slide, tile images centered on spot locations are selected (Fig. 1).

**Fig. 1.**
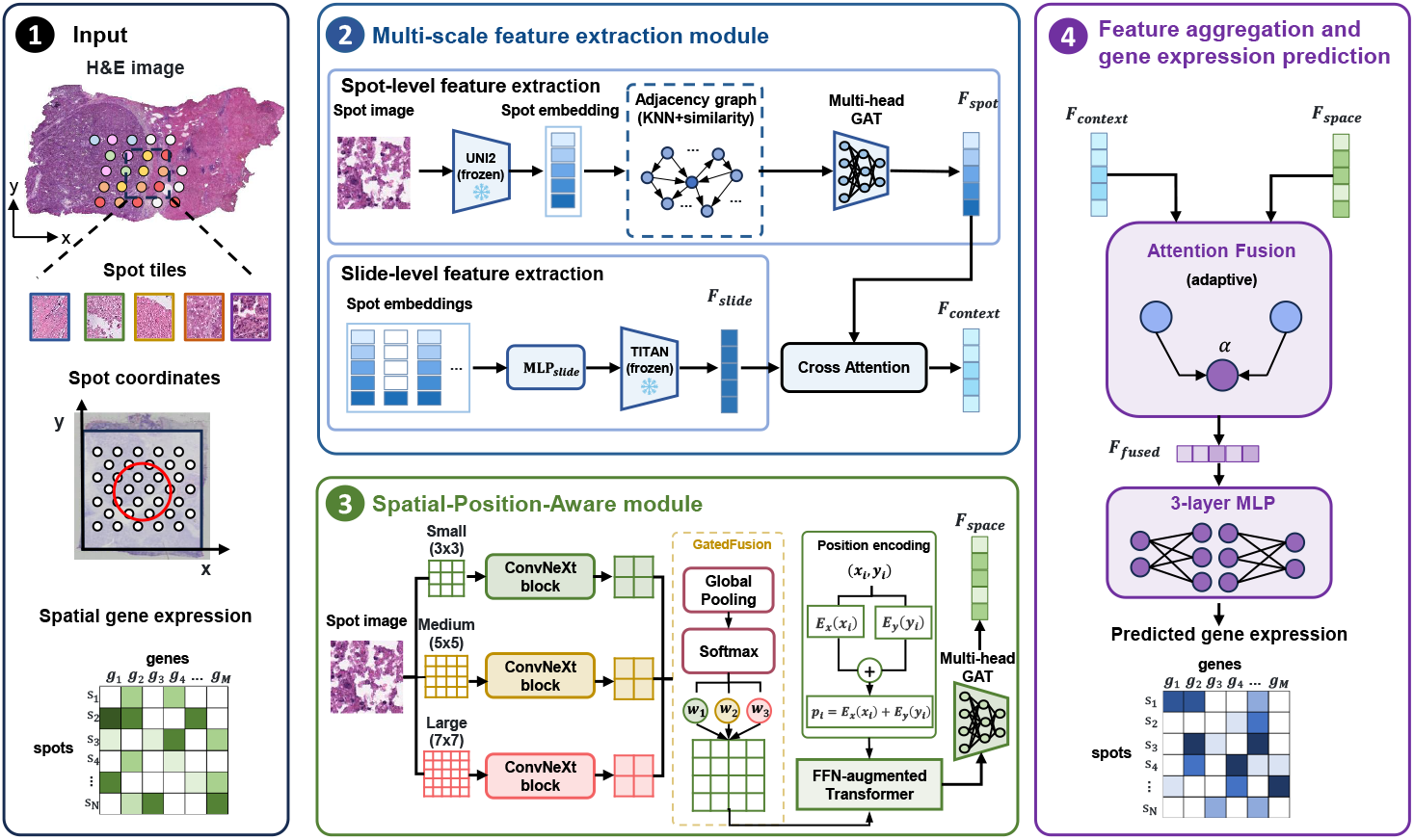
Overview of VISTA. For each tissue slide, spot-centered image tiles and spatial coordinates are extracted as model inputs to predict spatial gene expression. VISTA constructs a multi-scale semantic representation by combining spot-level features *F_spot_* and slide-level features *F_slide_*. A global-to-local cross-attention mechanism selectively incorporates slide-level context into spot-level embeddings to generate *F_context_*. In parallel, the Spatial-Position-Aware module extracts fine-grained local morphology using multi-receptive-field ConvNeXt branches with GatedFusion. To incorporate learnable coordinate embeddings, an FFN-augmented Transformer and multi-head GAT aggregation are applied to produce *F_space_*. Finally, features from the cross-scale semantic branch and the spatial-position-aware branch are adaptively fused via an attention mechanism. The resulting representation *F_fused_* is used by a three-layer MLP to predict spatial gene expression.

At the spot level, tile features were extracted using the UNI2 pathology foundation model [24], pretrained on over 200 million H&E images to provide biologically meaningful representations. A Multi-Head Graph Attention Network (GAT) [25] was then applied to aggregate information from neighboring spots. In parallel, the set of tiles in the whole slide was processed by the TITAN whole-slide foundation model [26] to extract a global feature that enhances spot representations. To bridge these two scale features, a global-to-local cross-attention mechanism selectively incorporates slide-level context into spot-level embeddings, thereby overcoming the locality limitation of tile-based representations.

To further complement semantic features from foundation models, we designed a Spatial-Position-Aware module to extract fine-grained information from spot images. Recognizing that each spot often contains genetic material from multiple cells with varying numbers, a multi-receptive field extractor based on ConvNeXt [27] blocks was employed. This module uses a GatedFusion mechanism to dynamically modulate the contributions of each convolutional branch, emphasizing the most informative spatial scale for each local region. To endow these features with spatial awareness and continuity, an FFN-augmented Transformer [12] and a secondary GAT were used to integrate the physical location embeddings of each spot.

Finally, features from the cross-scale semantic branch and the spatial-position-aware branch are adaptively fused via an attention mechanism, followed by a three-layer MLP to predict gene expression values at each spot. To address inter-sample variability arising from gene-level sequencing depth differences across slides [28], VISTA was trained with a composite loss function that jointly optimizes reconstruction and correlation-based regularization.

### 2.2 VISTA enables accurate prediction of spatial gene expression from H&E images

To evaluate the performance of VISTA in predicting spatial gene expression from H&E images, we conducted comprehensive benchmarking through both leave-one-section-out cross-validation and independent external validation. VISTA was compared against seven state-of-the-art methods: DeepPT [8], EGNv1 [14], TCGN [15], THi- toGene [16], FmH2ST [19], MISO [20], and HIST2ST [13]. Prediction performance was systematically evaluated at both the gene and spot levels using six quantitative metrics: Pearson Correlation Coefficient (PCC), Mutual Information (MI), Jensen–Shannon Divergence (JSD), Area Under the Curve (AUC), Concordance Index (C-index), and Structural Similarity Index (SSIM). To consolidate these metrics, we defined the Gene-wise Assessment Score (GAS) as the mean of all six metrics at the gene level. For spot-level evaluation, SSIM is inapplicable to individual spots [29], so we defined the Spot-wise Assessment Score (SAS) as the mean of the remaining five metrics. An overall performance score was then computed as the average of GAS and SAS, ensuring a balanced evaluation across molecular and spatial dimensions.

#### Leave-one-section-out cross-validation performance

Using a consistent training and evaluation protocol, we first performed leave-one-section-out cross-validation to benchmark our model against seven state-of-the-art models on the HER2+ breast cancer dataset [30], comprising 32 tissue sections from 8 patients. The models were iteratively trained on slides from all sections except one, and then tested on the held-out slides. VISTA demonstrated the highest performance across all evaluated metrics, outperforming existing methods in both gene-level and spot-level prediction while effectively capturing global expression profiles and local spatial heterogeneity (Fig. 2a).

**Fig. 2.**
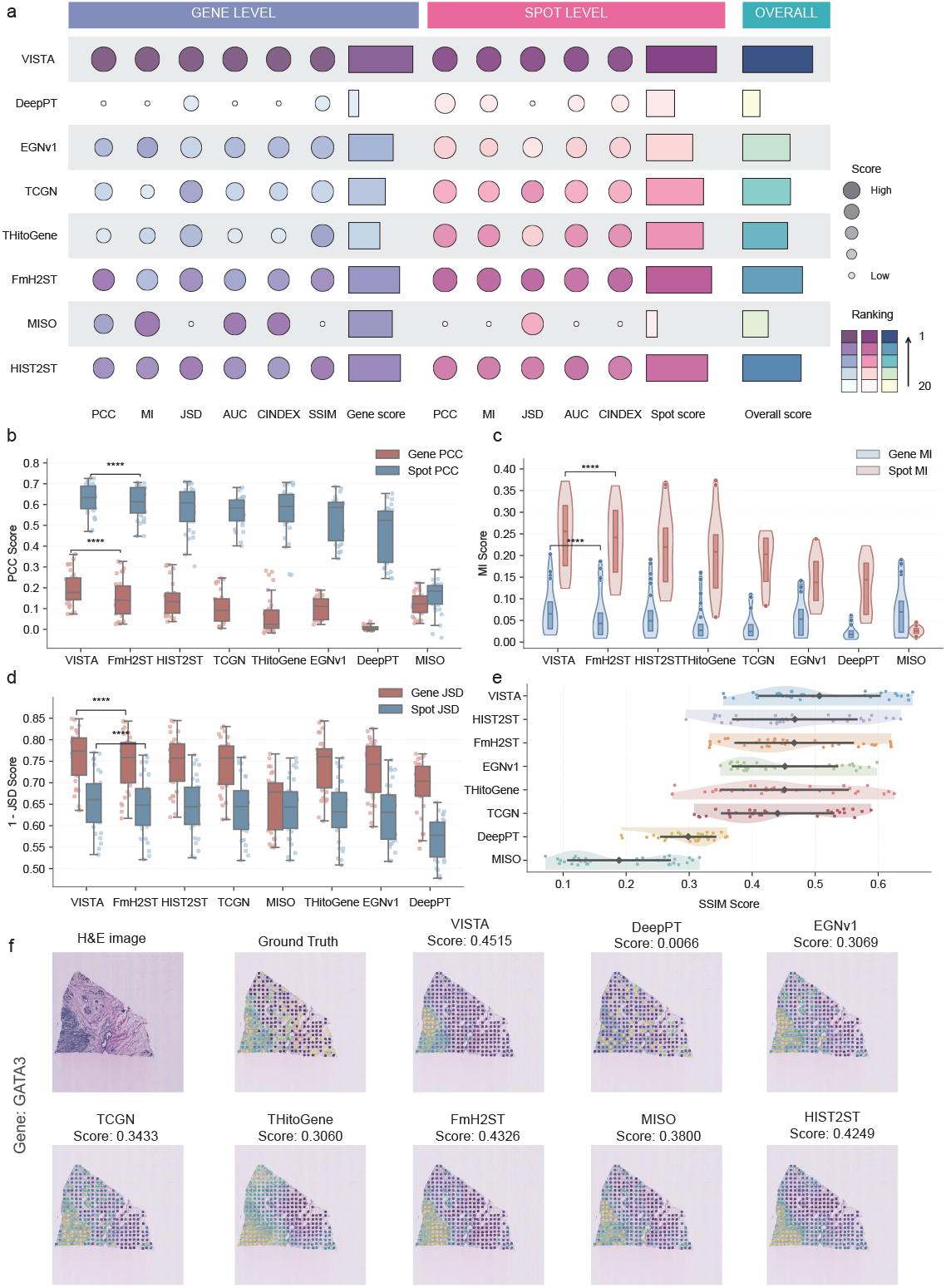
Leave-one-section-out cross-validation of VISTA for predicting spatial gene expression. **a,** Comprehensive comparison of VISTA with seven state-of-the-art methods, including DeepPT, EGNv1, TCGN, THitoGene, FmH2ST, MISO, and HIST2ST on the HER2+ breast cancer spatial transcriptomics dataset. Circle size represents the metric value, color indicates the relative ranking, and bar plots show the aggregated gene-level, spot-level, and overall scores. **b,** Box plots of gene-level and spot-level PCC scores between ground-truth and predicted gene expression. In the box plot, the center line denotes the median, and the box limits denote the upper and lower quartiles. Statistical significance determined by pairwise comparisons: **** *P* < 0.0001. **c,** Violin and box plots of gene-level and spot-level MI scores. **d,** Box plots of gene-level and spot-level 1-JSD scores. Higher 1-JSD values indicate better preservation of expression probability distributions. **e,** Horizontal violin and box plots of SSIM scores between ground-truth gene expression and predicted gene expression for each method. Overlaid points indicate individual test samples. **f,** Visualization of GATA3 expression in an example slide. Numbers above each prediction indicate the PCC score between the predicted and ground-truth GATA3 expression maps.

To evaluate predictive accuracy, we computed PCC between predicted and observed gene expression at both gene and spot levels. The gene-level and spot- level PCC averaged 0.196 and 0.629, surpassing the second-best FmH2ST (0.149 and 0.610), indicating the robustness of VISTA in predicting high-fidelity spatial transcriptomics (Fig. 2b). To directly evaluate how well predicted gene expression preserves the underlying distributional patterns, we computed 1-JSD between the probability distributions of predicted and observed expression. VISTA achieved the highest median 1-JSD across all compared methods, indicating that the predicted expression profiles conform to the original biological distributions (Fig. 2d). Beyond distributional similarity, SSIM was used to evaluate the spatial structural fidelity of predicted gene expression. VISTA achieved the highest SSIM score of 0.505, exceeding HIST2ST by 7.91%, indicating that its predictions maintain both structural integrity and spatial gradients (Fig. 2e). In terms of MI, AUC, and C-index, VISTA exhibited more stable and consistent predictions across sections, suggesting the predicted expression profiles aligned with the ground-truth and enabled better distinguishing between zero and non-zero expressions than other methods (Fig. 2c and Sup. Fig. 1).

To intuitively illustrate predictive performance, we visualized the expression pattern of GATA3 [31], a reliable breast tumor marker, on a slide (Fig. 2f). The result shows VISTA reappears with sharp spatial boundaries and fine-grained spatial patterns consistent with ground-truth, whereas competing methods often produce over-smoothed patterns. In contrast, competing methods, such as FmH2ST and HIST2ST, partially capture spatial heterogeneity but fail to fully preserve low-expression clusters, while methods like DeepPT and MISO show weaker correlation with the ground-truth. These results collectively demonstrate that VISTA not only quantitatively outperforms existing methods but also preserves biologically meaningful spatial structure, supporting accurate prediction of spatial gene expression patterns from H&E morphology.

#### Independent validation performance

To test the generalizability of VISTA, we trained all models on the full HER2+ ST dataset and then assessed their performance on five independent breast cancer datasets obtained from 10X Genomics [32], focusing on the 1,000 highly variable genes shared across datasets. The training cohort comprises low-density sections with an average of 361 ± 157 spots (*mean ± SD*), while the independent validation cohorts feature high-density capture with an average of 3477 ± 1345 spots (*mean ± SD*). This discrepancy in spot density makes the cross-dataset task more challenging, requiring the model to learn generalizable morphology-to-expression relationships rather than simply memorizing characteristics of the training dataset.

Despite these challenges, VISTA maintained superior performance across nearly all evaluation metrics, achieving a SAS of 0.552 and a GAS of 0.374, outperforming all seven competing methods (Fig. 3a). Notably, VISTA sustained its top ranking in PCC, MI, and JSD, which were most reflective of biological molecular concordance (Fig. 3b,c,d and Sup. Fig. 2). This indicated that the model successfully internalized correspondences between histological morphology and transcriptomic features that transfer across diverse patient cohorts and sequencing platforms. While VISTA ranked fourth in SSIM (0.151), this modest reduction reflects sensitivity to pixel-level variations in dense sections rather than loss of spatial fidelity (Fig. 3e).

**Fig. 3.**
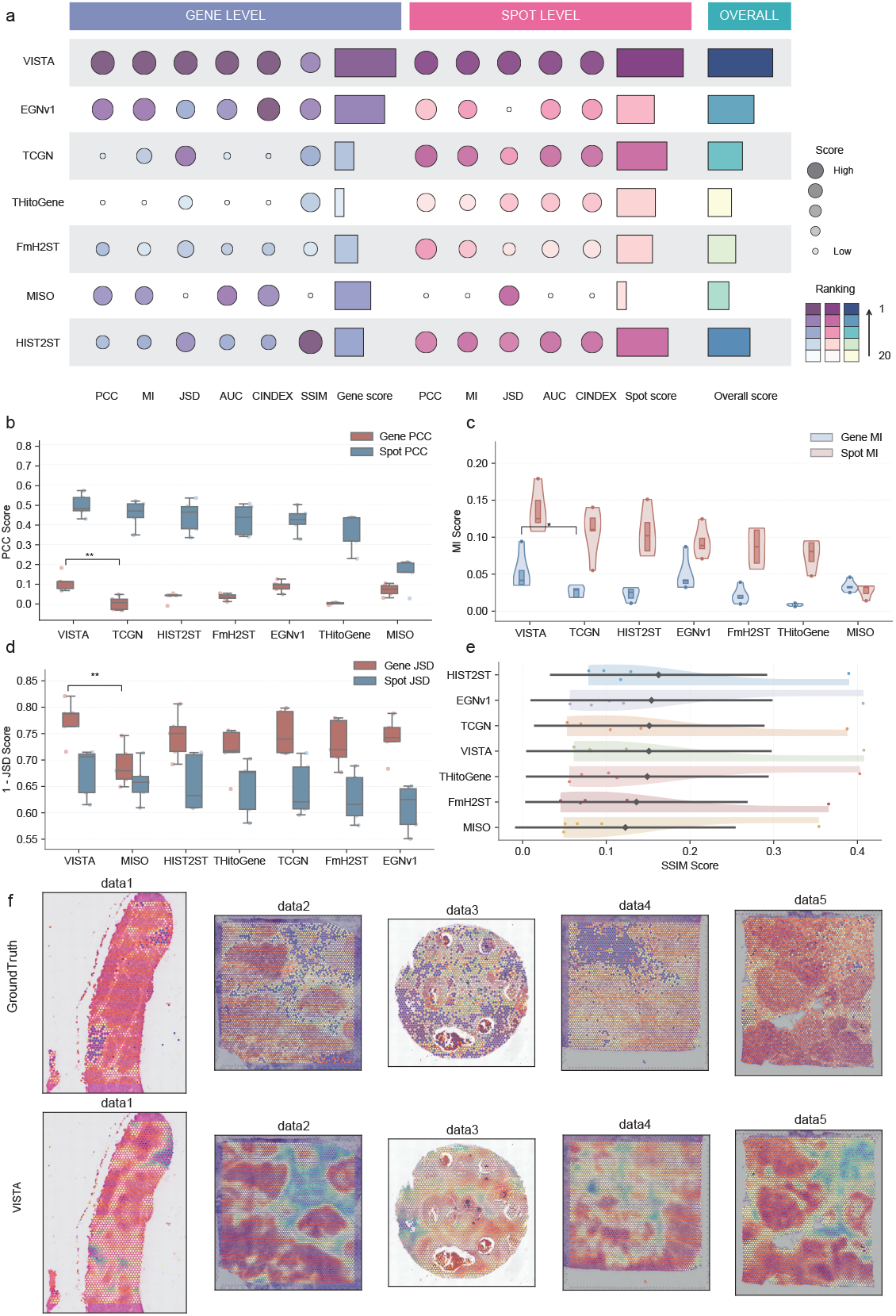
Independent validation of VISTA for predicting spatial gene expression. **a,** Comprehensive comparison of VISTA with competing methods, including EGNv1, TCGN, THitoGene, FmH2ST, MISO, and HIST2ST on the five independent breast cancer spatial transcriptomics datasets. Circle size represents the metric value, color indicates the relative ranking, and bar plots show the aggregated gene-level, spot-level, and overall scores. **b,** Box plots of gene-level and spot-level PCC scores between ground-truth and predicted gene expression. In the box plot, the center line denotes the median, and the box limits denote the upper and lower quartiles. Statistical significance determined by pairwise comparisons: **** *P* < 0.0001. **c,** Violin and box plots of gene-level and spot-level MI scores. **d,** Box plots of gene-level and spot-level 1-JSD scores. Higher 1-JSD values indicate better preservation of expression probability distributions. **e,** Horizontal violin and box plots of SSIM scores between ground-truth gene expression and predicted gene expression for each method. Overlaid points indicate individual test samples. **f,** Visualization of GATA3 expression across the five independent validation datasets. Ground-truth GATA3 expression maps are shown in the top, and VISTA-predicted expression maps are shown in the bottom.

Spatial visualization of GATA3 expression patterns across the independent validation cohorts further substantiates the superior generalization capacity of VISTA (Fig. 3f and Sup. Fig. 3). Compared to the Leave-one-section-out cross-validation, independent datasets present greater spatial complexity and expression heterogeneity. Despite the substantially increased spot density and spatial scale, VISTA successfully predicts tumor–epithelium boundaries and captures the enrichment of GATA3 within tumor regions. In contrast, methods that achieve competitive performance in cross-validation, such as FmH2ST and TCGN, show performance degradation when applied to high-density datasets. This ability confirms that VISTA achieves cross-dataset transfer through multi-scale feature integration and spatial awareness, whereas competing methods rely on dataset-specific optimization. Overall, the results of VISTA from both within-cohort cross-validation and cross-cohort independent validation demonstrate that it enables reliable spatial transcriptomics prediction across diverse histopathology collections.

### 2.3 Spatial transcriptomics prediction in TCGA identifies prognostic patient stratification and survival-associated domains

To evaluate whether spatial gene expression inference enables clinically meaningful prognostic stratification, we applied VISTA to 79 breast cancer samples from The Cancer Genome Atlas (TCGA) [33] with matched diagnostic H&E histopathological slides. Using the model pretrained on the HER2+ ST cohort, spatial expression profiles were predicted for each sample and aggregated into bulk-level transcriptomes for comparison with reference TCGA RNA-seq data. We first assessed the transcriptomic concordance between inferred expression profiles and TCGA bulk RNA-seq data. VISTA achieved a PCC of 0.48, ranking second among all compared methods and demonstrating that the inferred transcriptomic profiles retained agreement with reference bulk sequencing data (Fig. 4a). To evaluate prognostic relevance, we performed univariate Cox regression on the inferred expression profiles to identify survival-associated genes, and the top 5 genes were integrated into a multivariate Cox proportional hazards model. This yielded a C-index of 0.9 (1-SD = 0.94) with p = 0.002, outperforming all comparison methods. Kaplan-Meier survival analysis [34] further demonstrated that risk stratification based on predicted expression of VISTA clearly separated low and high risk patient groups (log-rank test, p = 0.002), with survival curves paralleled with the ground-truth (Fig. 4b and Sup. Fig. 4). To evaluate the biological relevance of inferred genes, we quantified the overlap between the top 100 survival-associated genes identified from predictions and those derived from TCGA bulk RNA-seq data. VISTA achieved the greatest overlap, indicating that the inferred gene expression preserves relevant prognostic signals more than competing methods (Fig. 4a). These results demonstrate that VISTA effectively translates histopathological information into prognostically meaningful molecular profiles.

**Fig. 4.**
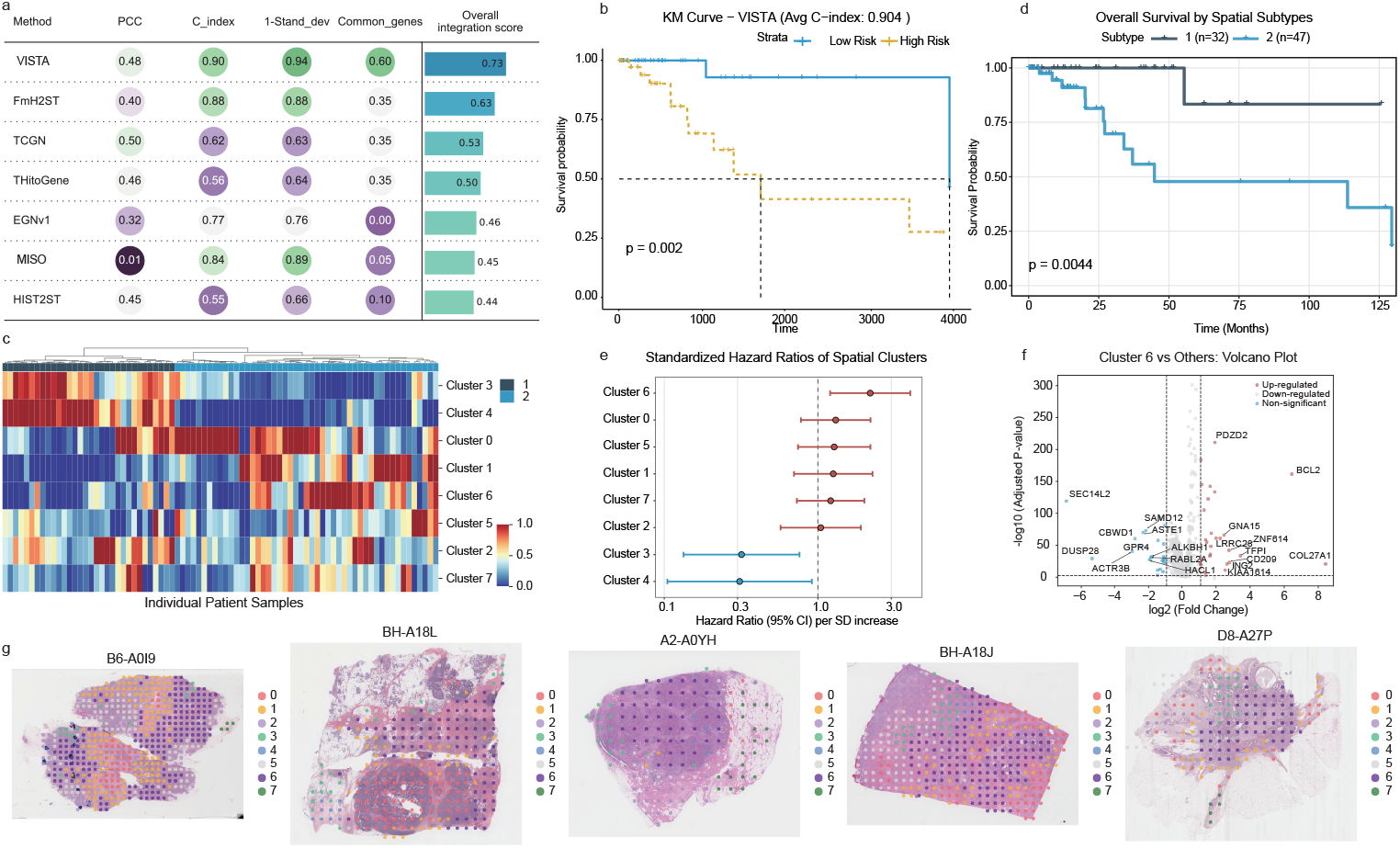
Identification of prognostic spatial subtypes and survival-associated spatial domains in the TCGA breast cancer cohort. **a,** Predicted spatial expression profiles were aggregated into pseudobulk transcriptomes and evaluated by PCC, C-index, model stability, and overlap with survival-associated genes identified from TCGA RNA-seq data. The overall score summarizes the four criteria. **b,** Kaplan-Meier survival curves for patients stratified according to the median risk score from the multivariate Cox model based on VISTA-inferred expression profiles. The average C-index is shown in the title, and the P value was calculated using a two-sided log-rank test. **c,** Hierarchical clustering of spatial cluster compositions for each tumor slide reveals two distinct spatial subtypes across n=79 TCGA breast cancer samples. The heatmap displays clusters in rows and patients in columns, with values indicating the proportional abundance of each cluster. The top annotation bar denotes the two spatial subtypes. **d,** Kaplan-Meier survival analysis comparing the two spatial subtypes identified from spatial cluster compositions. The P value was calculated using a two-sided log-rank test. **e,** Forest plot of standardized hazard ratios for the eight spatial clusters. Hazard ratios were estimated by univariate Cox regression using Z-score-normalized cluster proportions. Points indicate hazard ratios per standard deviation increase, horizontal bars indicate 95% confidence intervals, and the dashed vertical line marks HR = 1. **f,** Volcano plot showing differential gene expression between Cluster 6 and all other spatial clusters. The x-axis shows *log*_2_ fold change, and the y-axis shows *−* log_10_ adjusted P value. Red and blue points indicate upregulated and downregulated genes, respectively, and gray points indicate non-significant genes. Representative differentially expressed genes are labeled. **g,** Spatial visualization of the eight aligned spatial clusters overlaid on H&E images from representative TCGA samples, including B6-A0I9, BH-A18L, A2-A0YH, BH-A18J, and D8-A27P. Colors indicate spatial cluster identities, illustrating the tissue-level localization of survival-associated domains.

Having established gene-level prognostic fidelity, we next investigated whether incorporating spatial information could further refine patient stratification. SpaGCN was applied to identify spatial domains with coherent gene expression and histological context by aggregating gene expression of each spot from its neighboring spots [35]. To enable consistent cluster labeling across patients, the Louvain clustering algorithm [36] grouped all domains into 8 spatial clusters across the cohort, and each sample was represented by its cluster proportions (Fig. 4c). After hierarchical clustering, all samples were stratified into two distinct spatial subtypes, with each subtype characterized by similar spatial cluster compositions (Subtype 1: n=32, Subtype 2: n=47). Kaplan- Meier survival analysis revealed significant prognostic separation between these spatial subtypes (log-rank test, p = 0.0044), with Subtype 2 displaying marked reduced survival compared with Subtype 1 (Fig. 4d).

To determine which spatial clusters contributed to prognostic stratification, univariate Cox regression analysis was performed on the proportional abundance of each spatial cluster after Z-score normalization (Fig. 4e). Cluster 6 emerged as the sole significant survival domain associated with poor prognosis (HR = 2.19, 95% CI = 1.20–3.99, p = 0.011), with increasing Cluster 6 proportion linked to worse patient outcomes. Differential expression analysis of Cluster 6 versus other spatial clusters revealed a distinct phenotype associated with poor prognosis (Fig. 4f). Among the upregulated genes, COL27A1, BCL2, and CD209 were enriched for extracellular matrix remodeling and immune-associated regulation [37–39]. In contrast, downregulated genes, including SEC14L2, GPR4, and ALKBH1, were suggested to impair metabolic regulation and DNA/epigenetic repair-related processes [40–42]. These results suggest that Cluster 6 represents a tumor-associated spatial state characterized by stromal remodeling, immune-related alterations, and metabolic dysregulation. Spatial visualization of Cluster 6 across representative samples (B6-A0I9, BH-A18L, A2-A0YH, BH-A18J, D8-A27P) provides visual evidence for its prognostic significance (Fig. 4g). The visualization shows that Cluster 6 localized to tumor-enriched regions, segregating from surrounding stromal areas, thereby linking spatial tumor architecture to adverse prognosis. These results demonstrate that spatial gene expression inferred from VISTA enables meaningful patient stratification and identifies high-risk tumor-associated domains within the breast cancer microenvironment.

## 3 Discussion

Spatial transcriptomics revolutionized our comprehension of cellular heterogeneity, intercellular communication, and tissue structure by integrating spatial context with gene expression profiles [43]. However, the high cost of these spatial transcriptomics technologies has prevented their adoption in clinical and large-cohort settings. Predicting spatial gene expression from H&E images can enrich large gene expression profile datasets, facilitating comprehensive analysis and discoveries. In this study, we introduced VISTA to predict virtual spatial transcriptomic profiles from H&E images by integrating multi-scale histological representations and spatial awareness. This strategy enables predicted expression profiles to retain spatial information, providing a foundation for downstream analyses in pathology large-cohort.

Existing patch-based methods, such as DeepPT [8] and EGNv1 [14], primarily operate on isolated image patches, which assume that the gene expression of a spot can be inferred from its local appearance. This assumption neglects the fact that gene expression is influenced by neighboring cells and tissue boundaries that extend beyond the bounds of a single spot. VISTA addresses this limitation through its cross-scale fusion architecture, which integrates local tile-level features with global slide-level context to expand the receptive field and model inter-spot relationships. Although some existing methods, such as TCGN [15], THItoGene [16], and FmH2ST [19], incorporate neighborhood aggregation or graph attention to capture local spatial dependencies, they often overlook intra-spot heterogeneity and tissue architecture. To overcome this, VISTA employs a Spatial-Position-Aware module with GatedFusion, which adaptively integrates spatial coordinates and multi-scale visual features, preserving local structural variation and enhancing spot predictions. Furthermore, several emerging methods, such as MISO [20], leverage pretrained foundation models to improve feature extraction, which is still limited across scales. VISTA combines foundation model representations with cross-scale fusion, enabling the effective capture of both local detail and global organization. This dual-scale strategy reflects a key principle in histopathology practice that accurate diagnosis requires simultaneous assessment of local cellular structures and global tissue architecture [44].

Importantly, our work suggests the persistent performance advantage of VISTA across both cross-validation and independent external validation, particularly in high-density datasets where competing methods tend to over-smooth spatial expression patterns. By inferring spatial expression in TCGA breast cancer cohorts, we accurately stratified patients prognostically and predicted treatment response, highlighting spatial biomarkers associated with treatment outcomes. Furthermore, in liver cancer samples, VISTA revealed tissue heterogeneity and identified spatial domains enriched for distinct functional pathways, providing insight into tumor microenvironment organization. Through applying VISTA to a trastuzumab response cohort, spatial immune and signaling markers associated with treatment response were revealed, highlighting the spatial organization of the tumor microenvironment and improving the prediction of patient outcomes from H&E pathology slides. Overall, VISTA integrates multi-scale visual features and spatially aware mechanisms to link routine pathology images with spatial transcriptomics, enabling the extraction of molecular information from large clinical datasets without costly experiments and offering new insights into disease mechanisms and treatment response.

## 4 Methods

### 4.1 Preprocessing of spatial transcriptomic data

For 10x Visium spatial transcriptomics datasets, raw gene expression matrices were obtained from the Space Ranger output. To mitigate variability in sequencing depth across spots, we first performed library-size normalization, scaling each spot to a total count of 10,000. Subsequently, a *log*(*x* + 1) transformation was applied to stabilize variance and reduce the influence of highly expressed genes. Detailed statistics of the datasets, including sample size, number of spots, genes, and sparsity levels, are summarized in Sup. Table 1.

### 4.2 Preprocessing of histological images

Histological images from multiple cohorts were processed according to their respective platforms and imaging characteristics. For the HER2+ human breast cancer dataset acquired with the 10x Visium platform, the whole-slide image was divided into non-overlapping 224×224 pixel tile images centered around each spatial transcriptomics spot.

For the independent breast cancer datasets [32], the primary liver cancer atlas [45], and our in-house liver cancer cohort, the crop size was determined by the spot diameter fullres parameter provided in the spatial metadata, ensuring spatial alignment between each image tile and its corresponding gene-expression profile.

For clinical cohorts without paired spatial transcriptomics data, including TCGA-BRCA [33] and the Yale trastuzumab response cohort [46], whole-slide images in SVS format were converted to JPEG images and tiled into non-overlapping virtual spots. Each virtual spot was represented by a 20×20 pixel image tile, and the tiles composed of a whitish background were excluded by filtering out those whose mean RGB values exceeded predefined thresholds (*R >* 220, *G >* 220, *B >* 220). For each tile, both grid indices and pixel-level center coordinates were recorded to preserve spatial information for downstream graph construction and spatial transcriptomics inference.

### 4.3 Multi-scale feature extraction

Pathology foundation models pre-trained on large-scale histopathology collections can provide transferable morphological representations across diverse tissue types and disease categories [24, 47]. These models are valuable for spatial transcriptomics prediction, where paired image-expression training data are often limited. However, existing prediction methods mainly rely on features extracted from isolated tile images, neglecting the broader tissue architecture and slide-level context. To address this limitation, we designed a multi-scale framework consisting of spot-level and slide-level extraction modules. The two representations are subsequently combined through a global-to-local cross-attention mechanism.

#### 4.3.1 Spot-level feature extraction via UNI2

To obtain spot-level feature representations, we employed UNI2 [24], a vision transformer-based pathology foundation model pre-trained on over 100,000 whole-slide images using a combined DINOv2 [48] self-distillation and iBOT [49] masked image modeling objective. For a given H&E image, let 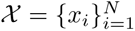 denote the set of image tiles centered at the spot coordinates, where *N* is the number of spots. Each tile was resized to 224 × 224 pixels and normalized according to the input requirements of UNI2 before being passed through the frozen encoder:

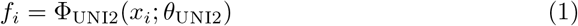

Where *θ*_UNI2_ denotes the frozen encoder parameters and *f_i_* represents the embedding of spot *i*. Although UNI2 produces high-quality tile embeddings, these representations are extracted independently for each spot and lack the spatial dependencies among neighboring spots.

To fully use spatial relationships between similar spots, we construct an undirected neighborhood graph *G* = (*V, E*), where each node *v_i_ ∈ V* is initialized with its corresponding tile embedding *f_i_*. Edge set *E*, and its weighted adjacency matrix *A* were constructed by integrating spatial distance with feature similarity. For each spot, we identified its *k* nearest spatial neighbors based on Euclidean distance computed from spatial coordinates. If spot *v_j_* is the neighbor of spot *v_i_*, then *A_ij_* = 1, otherwise *A_ij_* = 0. To ensure that edges reflect feature relevance, we modulated the spatial adjacency by the cosine similarity of the features, such that *A_ij_* = *β_ij_ ·* cos(*f_i_, f_j_*). A minimum node degree of *d*_min_ = 2 was enforced to maintain graph connectivity for spatially isolated spots.

Before graph-based aggregation, the UNI2 embeddings were projected into a 512- dimensional latent space through a two-layer MLP with SiLU [50] activation:

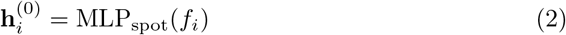

The projected spot embeddings were then passed to a Multi-Head Graph Attention Network (GAT) [25] to aggregate information from neighboring spots. For each connected pair (*v_i_, v_j_*), the unnormalized attention coefficien is computed as:

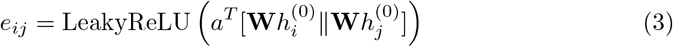

Where **W** is a shared linear transformation, *a* is a learnable attention vector, and denotes concatenation. The normalized attention coefficient was calculated over the neighborhood of spot *i*:

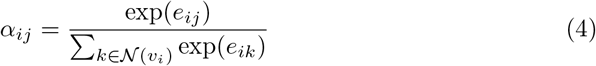

Each spot representation was updated by aggregating neighboring features weighted by the attention coefficients:

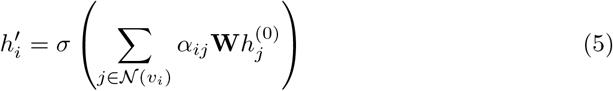

Where *σ*( ) denotes a nonlinear activation function. The outputs from all attention heads were concatenated and further aggregated to obtain the spot-level representation:

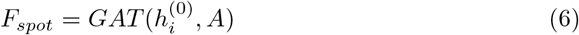

#### 4.3.2 Slide-level feature extraction

While spot-level embeddings capture local morphology, they may overlook global tissue organization and long-range contextual information. To encode slide-level context, we employed TITAN [26], a pathology foundation model designed for whole-slide representation learning. For a given section, the UNI2-derived spot embeddings *f_i_* were first projected into the feature dimension expected by TITAN using a two-layer MLP with SiLU activation:

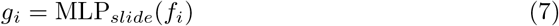

The projected embeddings, together with the pixel-level center coordinate **c***_i_*of the corresponding spots, were then provided to the frozen TITAN encoder:

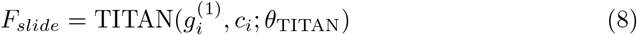

Where *θ*_TITAN_ denotes the frozen TITAN parameters. The resulting slide-level representation *F_slide_* anchors local spot predictions within the broader receptive field, resolving the spatial isolation limitation of traditional patch-based modeling. This slide-level representation provides global contextual information for cross-scale feature fusion.

#### 4.3.3 Global-to-Local feature fusion

To integrate local spot-level features with global slide-level features, we implemented a Global-to-Local Cross-Attention module. This module allows each spot to selectively incorporate information from the slide-level representation, expanding the receptive field beyond the local region. Specifically, *F_spot_*serves as the query, whereas *F_slide_*serves as the key and value, and cross-attention was computed as:

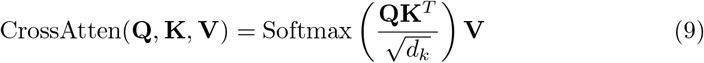

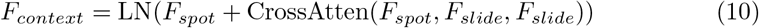

Where *d_k_*is the scaling factor corresponding to the feature dimension and LN(·) denotes layer normalization. The residual connection preserves the local specificity of each spot while the cross-attention pathway enriches it with global context.

### 4.4 Spatial-Position-Aware module

To complement the high-level semantic representations extracted by foundation models, we developed a Spatial-Position-Aware module to extract local morphology and spatial organization. This module extracts multiple receptive field morphological features from each spot image and injects coordinate information to encode both absolute tissue location and local neighborhood continuity. It provides spatially aware morphological features that complement the semantic context representation described above.

#### 4.4.1 Multi-receptive field feature extraction with GatedFusion

Because each spatial transcriptomics spot may cover multiple cells and heterogeneous local tissue structures [2], a single receptive field may be insufficient to capture the morphological information associated with spot-level gene expression. Therefore, we designed a multi-receptive-field convolutional extractor based on ConvNeXt [27] blocks. It can capture long-range morphological dependencies without sacrificing the necessary translation invariance. For each spot image tile *x_i_*, three parallel convolutional branches with different kernel sizes were applied to capture local patterns at small, medium, and large spatial scales:

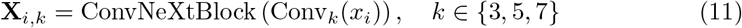

Where **X***_i,k_* denotes the feature map extracted from the branch with kernel size *k*. To adaptively modulate information from different spatial scales, we introduced a GatedFusion mechanism. The feature maps from the three branches were first summarized by global average pooling and concatenated. A two-layer gating network then produced normalized branch weights:

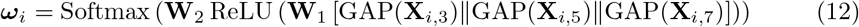

Where GAP(*·*) denotes global average pooling, denotes concatenation, **W**_1_ and **W**_2_ are learnable parameters, and ***ω****_i_* = *ω_i,_*_3_*, ω_i,_*_5_*, ω_i,_*_7_ represents the scale-specific gating weights for spot *i*. The fused multi-scale feature map was computed as:

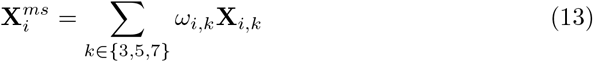

The fused feature map was further summarized into a spot-level morphological vector:

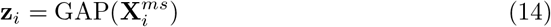

This adaptive fusion strategy allows the model to select the most informative receptive field for each local tissue region.

#### 4.4.2 Position-Aware spatial encoding via FFN-Transformer

To incorporate explicit spatial information, we encoded the physical coordinate of each spot into a learnable position embedding. Let **c***_i_* = (*u_i_, v_i_*) denote the pixel-level center coordinate of spot *i*. The coordinate embedding was defined as:

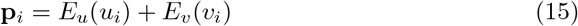

Where *E_u_*(*·*) and *E_v_*(*·*) are learnable embedding functions for the two coordinate axes. The position embedding *p_i_* was injected into the multi-scale morphological vector **z***_i_* through an FFN-augmented transformer-style block [12]. For the collection of spot-level features in a tissue section, we initialized:

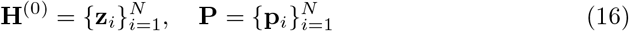

The position-aware feature representation was then updated as:

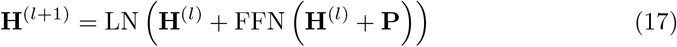

Where LN( ) denotes layer normalization, FFN(·) denotes a feed-forward network, and *l* indexes the layer. This formulation anchors each spot representation to its physical location within the tissue section while preserving the underlying morphological feature structure. Although coordinate injection encodes absolute spatial location, it does not explicitly model relative dependencies between neighboring spots. Therefore, the output of the final position-aware layer was further passed through a multi-head graph attention network:

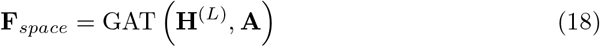

By combining coordinate-aware feature encoding with graph-based neighborhood aggregation, *F_space_*encodes both the physical location of each spot and its spatial relationships to the neighboring spots.

### 4.5 Feature aggregation and gene expression prediction

The context-enhanced representation *F_context_* from the multi-scale feature extraction module and the spatial-position-aware representation*F_space_* encode complementary information for each spot. To adaptively integrate these two feature sources, we used an attention-based fusion mechanism. For each spot *i*, the fused representation is:

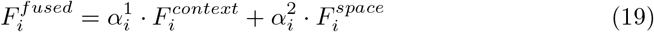

Where the attention weights 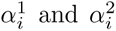 are computed by a learnable query-key

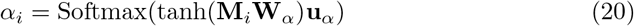

Where **M***_i_* is the concatenated modality matrix. The fused embedding 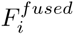 is finally input into a three-layer MLP to predict spatial gene expression. Detailed architectural specifications and training configurations are provided in Supplementary Note 1.

